# A hybrid approach for predicting multi-label subcellular localization of mRNA at genome scale

**DOI:** 10.1101/2023.01.17.524365

**Authors:** Shubham Choudhury, Nisha Bajiya, Sumeet Patiyal, Gajendra P. S. Raghava

## Abstract

In the past, number of methods have been developed for predicting single label subcellular localization of mRNA in a cell. Only limited methods had been built to predict multi-label subcellular localization of mRNA. Most of the existing methods are slow and cannot be implemented at transcriptome scale. In this study, a fast and reliable method had been developed for predicting multi-label subcellular localization of mRNA that can be implemented at genome scale. Firstly, deep learning method based on convolutional neural network method have been developed using one-hot encoding and attained an average AUROC - 0.584 (0.543 – 0.605). Secondly, machine learning based methods have been developed using mRNA sequence composition, our XGBoost classifier achieved an average AUROC - 0.709 (0.668 - 0.732). In addition to alignment free methods, we also developed alignment-based methods using similarity and motif search techniques. Finally, a hybrid technique has been developed that combine XGBoost models and motif-based searching and achieved an average AUROC 0.742 (0.708 - 0.816). Our method – MRSLpred, developed in this study is complementary to the existing method. One of the major advantages of our method over existing methods is its speed, it can scan all mRNA of a transcriptome in few hours. A publicly accessible webserver and a standalone tool has been developed to facilitate researchers (Webserver: https://webs.iiitd.edu.in/raghava/mrslpred/).

**Key Points:**

- Prediction of Subcellular localization of mRNA
- Classification of mRNA based on Motif and BLAST search
- Combination of alignment based and alignment free techniques
- A fast method for subcellular localization of mRNA
- A web server and standalone software

## Introduction

Messenger RNA (mRNA) is a single-stranded RNA, a transcription product that leads to protein synthesis via translation. It carries the cell’s genetic information from the nucleus to the cytoplasm. In the cytoplasm, mRNA is localized to different parts of the cell, resulting in an asymmetric distribution of proteins within the cell, thus establishing cell polarity [1]. It plays an important role in several developmental processes, such as neuronal maturation, embryonic patterning, cell migration, cell fate determination, cell adaptation to stress, and the development of body axes in Drosophila melanogaster [2–6]. Identifying the cellular location of mRNA provides valuable information about the amount of protein synthesis and the location, which correlates with its function [7,8]. While transporting mRNA over a protein has significant advantages, such as transportation costs reduction by the expression of mRNA to generate different types of localized proteins, rapid response to external stimuli, segregation of transcripts to specific organelles or compartments, and prevention of ectopic action of proteins during localization [1,8–10]. Therefore, knowing the subcellular localization of mRNA is important to understand various biological processes [8].

Studies on mRNA spatial localization within cells have been extensively studied recently. In-vitro visualization remains the gold standard for mRNA location to trace its subcellular localization. It was performed using improved classical in-situ hybridization such as fluorescent in-situ hybridization (FISH), MS2-system-based techniques, and high-throughput RNA sequencing with high sensitivity and selectivity [1,11,12]. Still, at the same time, they are laborious, limited applications to specific tissues, are expensive processes, and often require sophisticated instrumentation [13]. Advanced sequencing techniques generate a large amount of information about transcripts. Therefore, computational approaches are needed as alternative methods to determine the precise location of RNA transcripts in the cell at the level of the entire transcriptome [13].

Several databases have been created using the experimental data, which contain comprehensive information on the subcellular localization for all kinds of RNA, such as RNALocate and its updated versions [14,15], lncATLAS [16] and lncSLdb [17]. Novel computational RNA subcellular localization prediction tools developed from these databases, based on different machine learning methods as an efficient and effective approach, such as RNATracker [18], iLoc-mRNA [11], DM3Loc [1], mRNALocater [8], and mRNALoc [19]. Among tools using Cefra-Seq and Apex-RIP data to predict subcellular localization, RNAtracker [18] is a very popular tool. It deploys a deep recurrent neural network-based model to perform mRNA subcellular localization. However, the Cefra-Seq and APEX-RIP datasets are very noisy and are not reliable at times. Majority of tools have used localization data from RNALocate [15], which includes iLoc-mRNA [11], mRNALocater [8] and mRNALoc [19]. iLoc-mRNA [11] uses a SVM classifier to assign a single subcellular location to each mRNA sequence. It is designed to predict localization to four main locations - cytosol/cytoplasm, ribosome, endoplasmic reticulum and nucleus/exosome/dendrite/mitochondrion. mRNALoc [19] uses pseudo k-tuple nucleotide composition features to train a SVM classifier for subcellular localization prediction. It can assign one location out of five locations (Mitochondria, Cytoplasm, Nucleus, Endoplasmic reticulum, Extracellular region) to a mRNA sequence. mRNALocater [8] uses an ensemble learning algorithm by combining the outputs from three machine algorithms (LightGBM, XGBoost, CatBoost). The mRNA sequences are encoded into fixed length vectors by generating electron-ion interaction pseudopotential values and pseudo k-tuple nucleotide composition. The locations that the mRNALocater [8] can classify as same as mRNALoc [19]. These methods have been developed on an older version of RNALocate [14] and the updated version of RNALocate [15] has a greater number of mRNA sequences and low redundancy.

Lin et.al. have recently developed DM3Loc [1], which is a multi-label subcellular localization prediction tool that uses a deep learning framework. DM3Loc generates features by converting mRNA sequences into one-hot encoded vectors. These vectors are provided as input to a CNN classifier and uses a multi-head self-attention mechanism to improve its ability to identify sequence regions relevant to localization. However, using one-hot encoded RNA sequences has its own drawbacks, with the primary issue being the large size of input vector when we attempt to predict localization for a large number of mRNA sequences. This makes the tool inaccessible to researchers with minimal computational resources as frequent RAM crashes were observed for a larger query dataset. Also, computational time for a complex deep-learning model is considerably higher compared to a machine learning model.

Currently, most of the existing tools are single-label classifiers, i.e., they can classify mRNA into a single location. But in a realistic scenario majority of the mRNA travel throughout the cell and can be found at multiple locations within the cell. Also, tools designed based on deep learning models are very computationally intensive, and the standalone requires significant amount of resource to run. In this study, we have tried to address these issues by developing a simple multilabel classifier – MRSLpred, using a machine learning model that can identify all the possible locations of mRNA sequences based on compositional features, mimicking the ground reality.

## Material and methods

### Dataset collection

Pre-processed data was obtained from DM3Loc [1], where they have used both experimentally validated as well as database curated mRNA sequences of Homo sapiens from RNALocate database v2 [15]. It is a database dedicated to providing high confidence RNA subcellular localization information sourced from literature, other databases and RNA-seq datasets. Majority of the mRNAs were localized in more than one subcellular compartment which is generally the case with most mRNAs in a real-world scenario. Our dataset contains a total of 17,277 nonexclusive mRNAs and they are spread across six subcellular compartments: ribosome, cytosol, endoplasmic reticulum (ER), membrane, nucleus, and exosome. The dataset is graphically represented in Figure 1. The number of mRNAs with localization at different compartments are 11923 for nucleus, 17156 for exosome, 2338 for cytosol, 5210 for ribosome, 3232 for membrane and 1976 for ER. The distribution of location labels is depicted in Figure 2.

**Figure 1.**
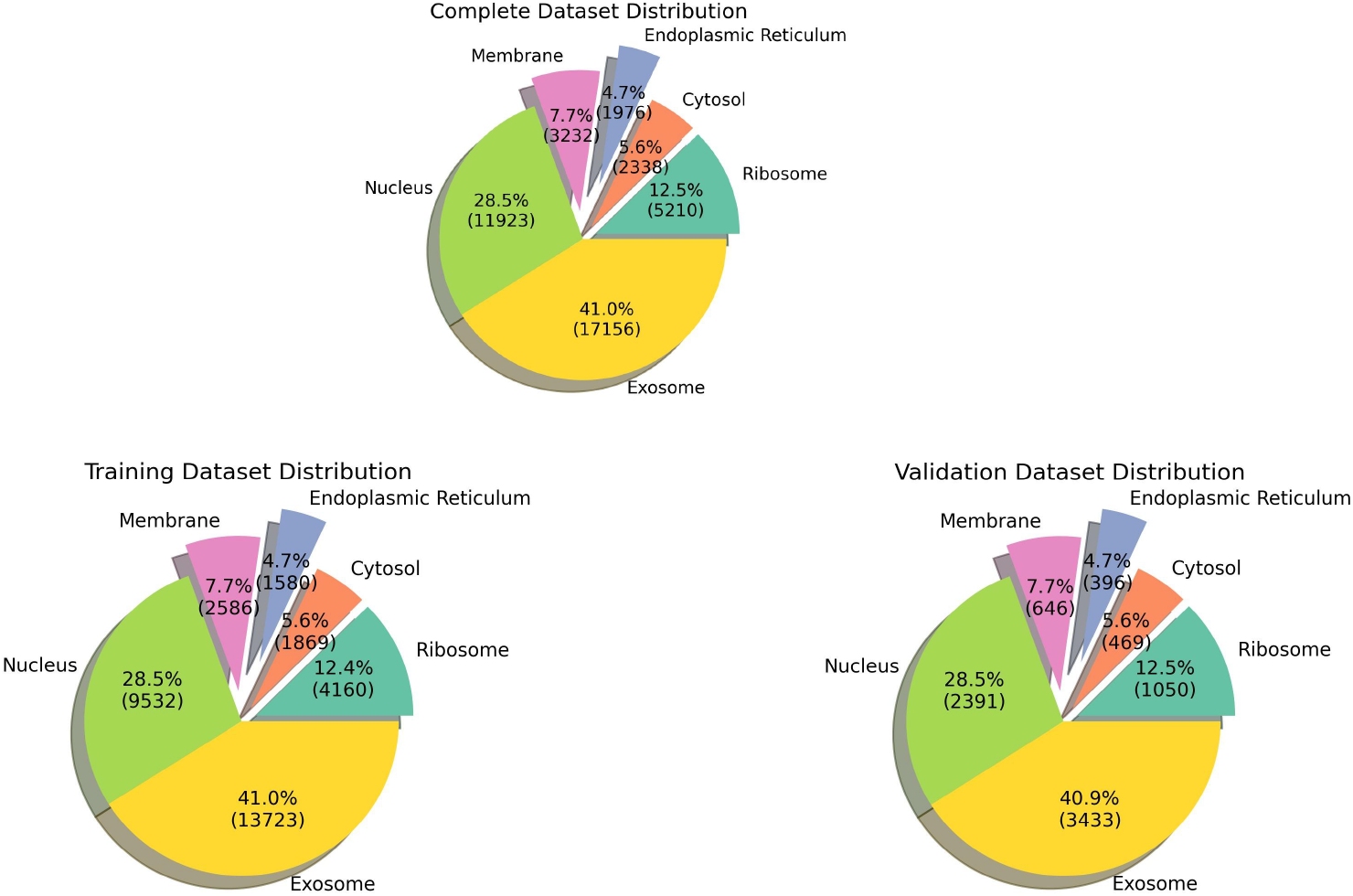
Pie Chart indicating the data distribution in all the datasets

**Figure 2.**
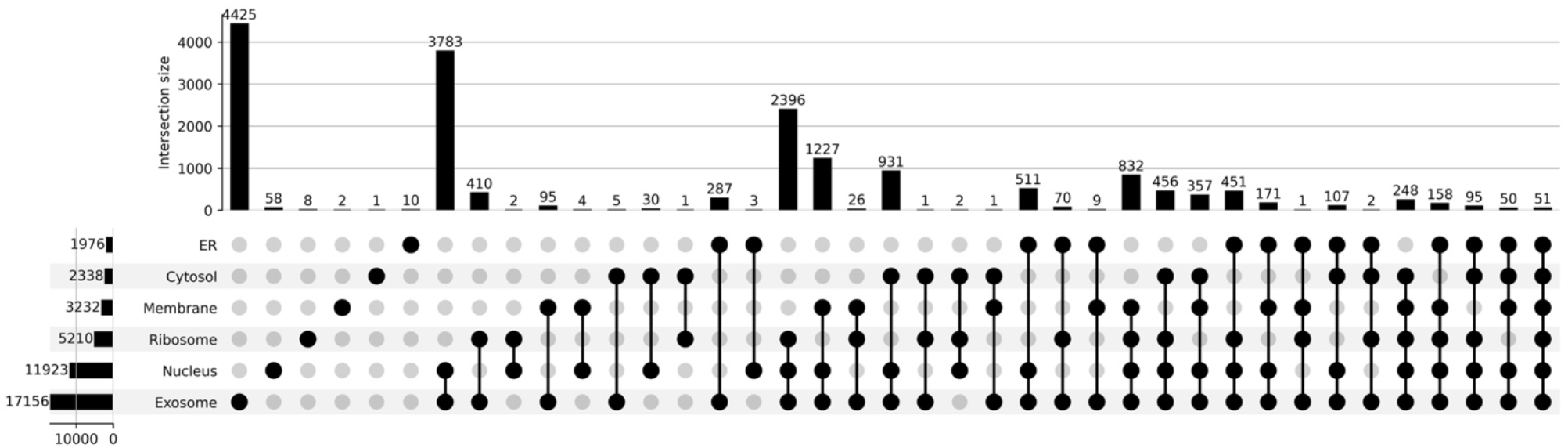
Upset plot depicting all the possible combinations of subcellular locations

### Feature generation

#### Composition-based features for machine learning models

In order to train our model, we are required to generate features or descriptors corresponding to each mRNA. For the aforementioned purpose, we have used the tool ‘Nfeature’ [20] [https://doi.org/10.1101/2021.12.14.472723], which can generate hundreds of features for a single mRNA sequence. These are the two feature classes which were used for training the models:

1. Composition of DNA/RNA for k-mer (CDK): k-mers of length 3 were generated by Nfeature and the frequency of each k-mer was used as a feature for training the ML model. It was calculated using the following formula

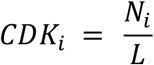 Where *N*_i_ represents the number of occurrences of the k-mer *i* in the mRNA sequence and *L* represents the length of the mRNA. For example, if ‘ATG’ is a 3-mer, the program will count all the instances of this 3-mer in the input sequence and divide it by the length of the input sequence.
2. Reverse complement of DNA for k-mers (RDK): k-mers of length 4 were generated by Nfeature and the frequency of the reverse complement of this k-mer will be used as a feature. The formula used for calculating RDK is similar to the one used for CDK. However, in this case instead of counting the occurrence of the k-mer in the input sequence directly, first the reverse complement of the sequence is generated and then the 4-mer frequency in that sequence is calculated.

Both these features were combined together to obtain a vector of 200 features for each mRNA.

#### One-hot encoding of mRNA sequences

Initially a CNN model was trained using one-hot encoded mRNA sequences. One-hot encoding was done by converting every mRNA sequence into a 2000 × 4 matrix where the columns represent the four nucleotides (A, T, C, G) and the rows represent the sequence information. For sequences greater than 2000 base pairs (bp), 1000 bp were taken from the 5’-end and the remaining 1000 bp were taken from the 3’-end. The sequences which have less than 2000 bp were used as it is and the remaining matrix was filled with zeros for all the nucleotides.

### Alignment-based methods

#### Similarity search using BLAST

In this study, we have used BLAST for determining sequence similarity, as it is the gold standard tool for finding similarity between sequences. Here, we have used it for assigning labels to mRNA sequences based on their similarity with sequences present in the database. The training dataset was used as a database, created using the makeblastdb binary within NCBI-BLAST version 2.13.0+. Subsequently the sequences of the validation dataset were queried against the database, using the blastn tool. The labels of the top hit (one with lowest e-value) for each query were then assigned to the query sequence. In order to identify the optimum e-value for running blastn, we have tried e-value cut-offs ranging from 10^-6^ to 10^3^ in increasing powers of 10.

#### Motif search using MERCI

Motifs are known to play a significant role in the localization of mRNA within the cell. We have used MERCI tool [21] to identify the presence of conserved motifs in the training dataset. For each location, the dataset was split into a positive and negative dataset based on the localization in that location. Both the positive and negative datasets are then provided as inputs to MERCI [21] and then the tool identifies motifs that can discriminate between positive and negative samples for that location. We acquired six sets of motifs specific to each location and this information was used to modify the probabilities for each location. For instance, in case a motif specific to ribosome sequences is found within a query sequence, the probability (provided by the ML model) that the query sequence belongs to ribosome, is updated to 1. So, the presence of motifs will practically override the prediction made by the ML model and for those sequences which do not have any motifs, remain unaffected.

#### Alignment-free methods – Machine learning and Deep learning models

The location label of each mRNA was generated using one hot encoding by converting locations into 0s or 1s, i.e., if a mRNA is only present in ribosome and nucleus, it will have a label like [1,0,0,0,1,0]. Initially a CNN model was trained using one-hot encoded mRNA sequences. One-hot encoding was done by converting every mRNA sequence into a 2000 × 4 matrix where the columns represent the four nucleotides (A, T, C, G) and the rows represent the sequence information. For sequences greater than 2000 base pairs (bp), 1000 bp were taken from the 5’-end and the remaining 1000 bp were taken from the 3’-end. The sequences which have less than 2000 bp were used as it is and the remaining matrix was filled with zeros for all the remaining positions. Composition-based features defined above were used to train various machine learning models. Model training was done on python using standard machine learning functions from scikit-learn and xgboost. Number of machine learning approaches were used to construct prediction models like Logistic Regression, Decision Tree, Random Forest Classifier, MLP Classifier, AdaBoost Classifier, Gaussian Naïve Bayes, Quadratic Discriminant Analysis, Gradient Boosting Classifier and Xtreme Gradient Boosting Classifier. The scikit-learn python library was used to implement these classification approaches.

Training of the models was done using ML classifiers combined with a multioutput classifier. The specialty of this model is that it takes CDK and RDK as features and predicts all the possible locations of the mRNA in one single step.

#### Five-fold cross validation

To ensure proper fitting of our model, we used five-fold cross validation (for both ML and DL models) to train and validate the models. The entire dataset was split in an 80:20 ratio where 80% of the data was used for training and 20% data was used for validation. The training data was further split into five parts and five-fold cross validation was performed on the same. In each iteration, a different fold was used for validation and the remaining four folds were used for validation. The training performance is calculated by taking the average over five iterations. The splitting of data was done in a stratified manner, ensuring that all the locations are equally distributed within each of the folds. Once the training was done, the model was validated using the 20% validation dataset.

#### Performance metrics

Evaluation of the model performance was done using standard performance metrics. The performance metrics used were - Sensitivity, Specificity, Accuracy, Matthews’ Correlation Coefficient, F1-score and Area Under the Receiver Operator Characteristics (AUROC). Out of these metrics, only AUROC is threshold independent, whereas all the remaining metrics are dependent on the threshold cut-off. The cut-off for the probabilities was determined by balancing out the sensitivity and specificity.

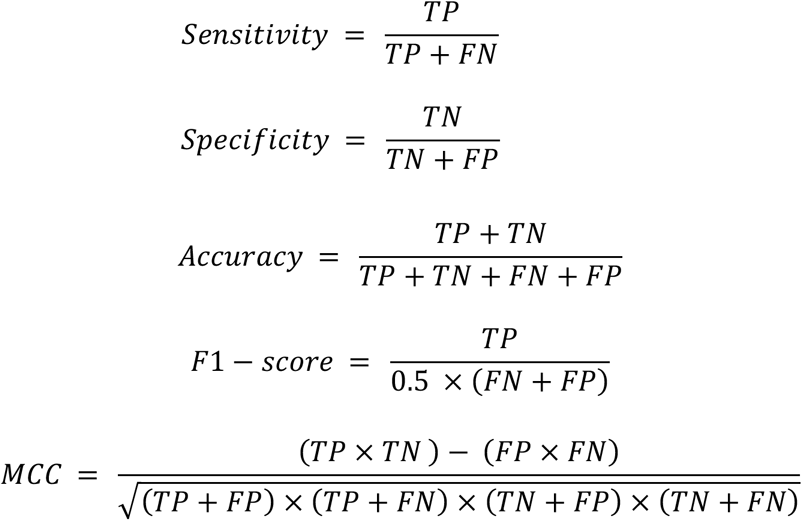

where TP is True Positive, FP is False Positive, FN is False Negative and TN is True Negative.

## Results

In this study, we have used a total of 17,277 mRNA sequences with non-exclusive locations for training our model. The primary objective is to develop a model that can accurately predict multiple locations for a single mRNA, mimicking a practical scenario within the cell. So, we have developed a multilabel classifier to predict subcellular localization of mRNA. The outline of the study is explained in Figure 3.

**Figure 3.**
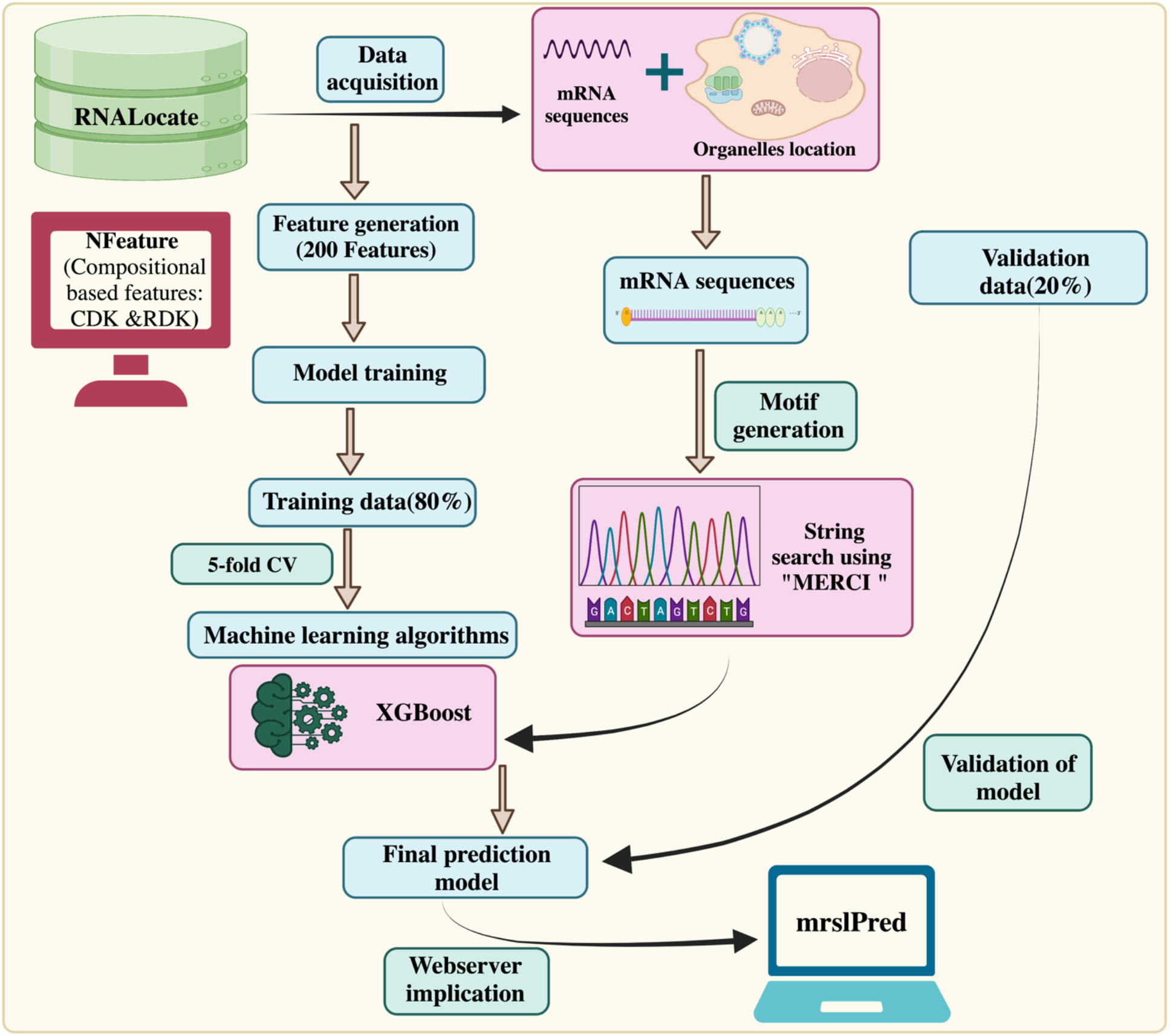
Outline of the methodology followed by MRSLpred

### Alignment-based methods

#### Performance of BLAST-search module

For assigning labels to mRNA sequences, we have used a similarity search method using BLAST. In this module, we have used BLAST to query the mRNA sequences of the validation dataset against the database created from the training dataset. The location labels for the top hit for each query mRNA were assigned to that mRNA. Different values of e-values were used to identify the optimal e-value where the BLAST module performed the best. Results for e-value = 10^-6^ are attached below (Table 1) –

**Table 1.** Search hits for BLAST module at e-value=10^-6^ (Total no. of hits = 2366)

|  | # of correct hits | # of incorrect hits |
| --- | --- | --- |
| Ribosome | 1532 | 834 |
| Cytosol | 1720 | 646 |
| ER | 1912 | 454 |
| Membrane | 1711 | 655 |
| Nucleus | 1602 | 764 |
| Exosome | 2338 | 28 |

It was observed that, at higher e-values the BLAST module performs poorly, as BLAST will allow random hits at higher e-values (Table 2). So, even though the number of hits increases at higher e-values but the performance decreases.

**Table 2.**
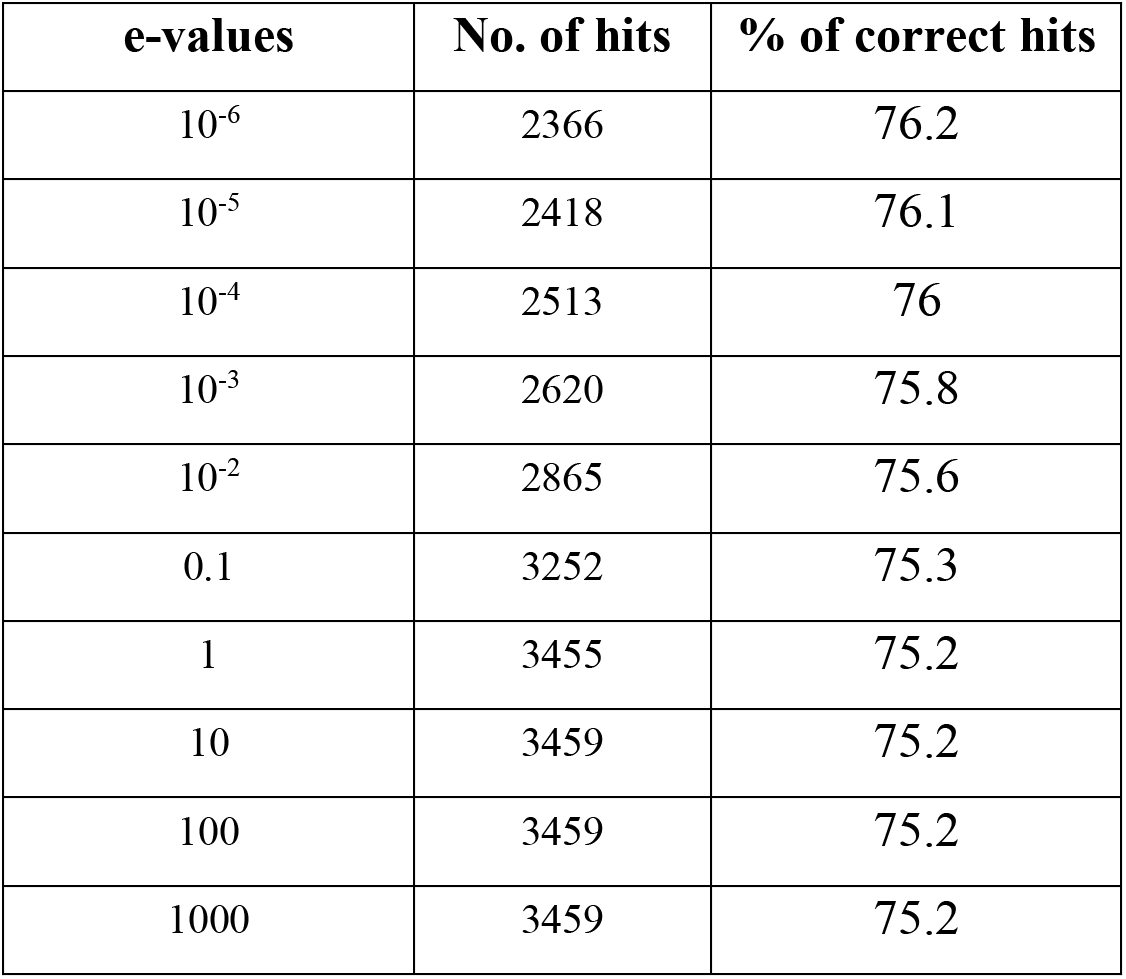
Comparison of performance among BLAST modules at different e-values

#### Performance of Motif-search module

Nucleotide motifs are known to affect mRNA localization and, in this study, we have tried to implement this ideology. Discriminatory motifs for each location were identified using the training dataset and presence of these motifs were then used to assign location labels to each mRNA in the validation dataset. The motifs, that are unique to the positive dataset for each location, are searched in the validation dataset and if any one of these motifs are found within the sequence, then the label for that mRNA in that location is assigned as 1 or 0 otherwise. The number of hits for motifs in each individual location were as follows – Ribosome: 33, Cytosol: 25, Endoplasmic Reticulum: 66, Membrane: 29, Nucleus: 96 & Exosome: 1655. Majority of the motifs were predominantly identified in the mRNA sequences that were assigned to exosomes. It may be possibly due to the relatively high number of sequences assigned to the exosome location.

The performance metrics for the prediction done by motif module are provided in Table 3.

**Table 3.**
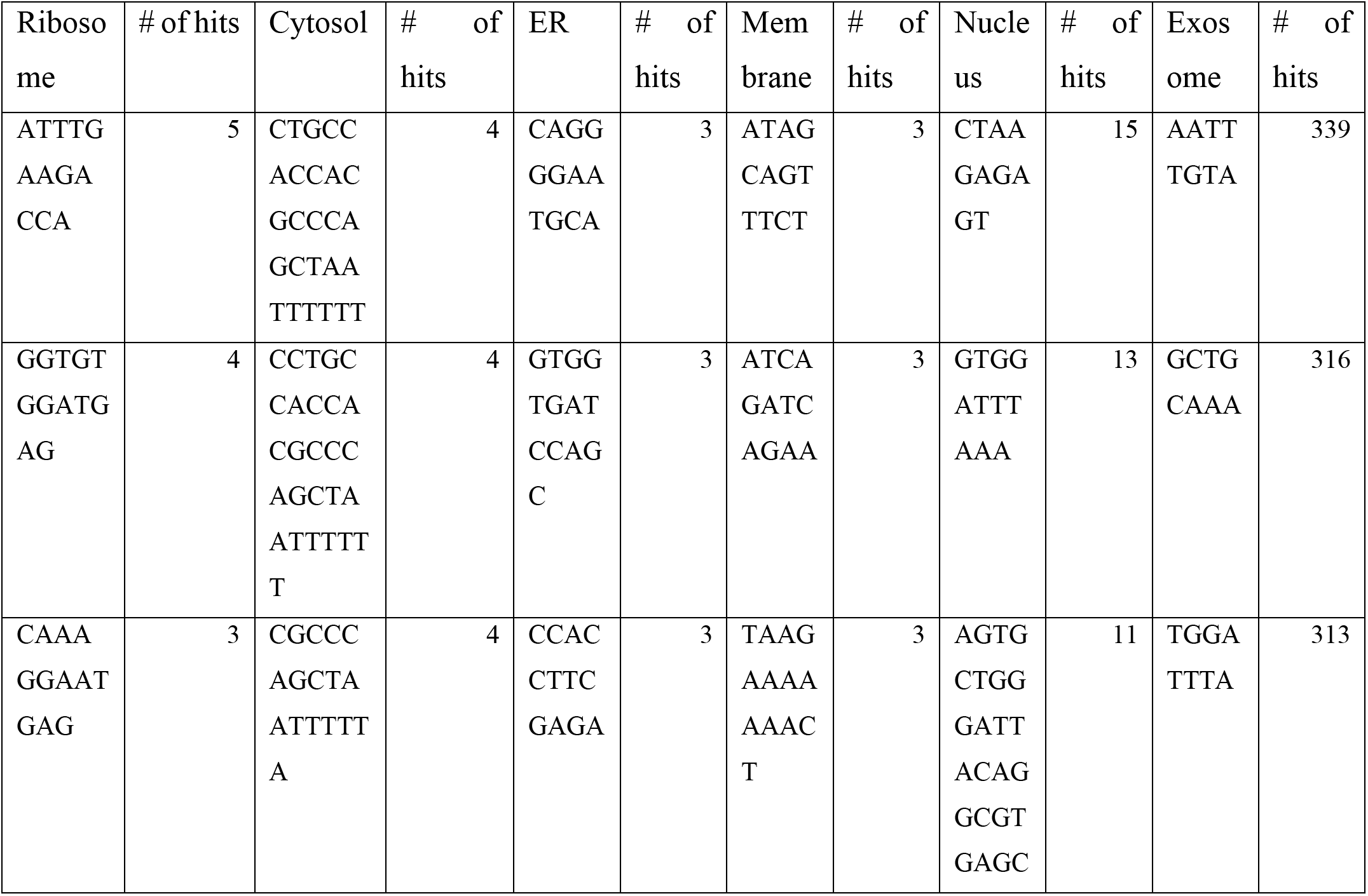
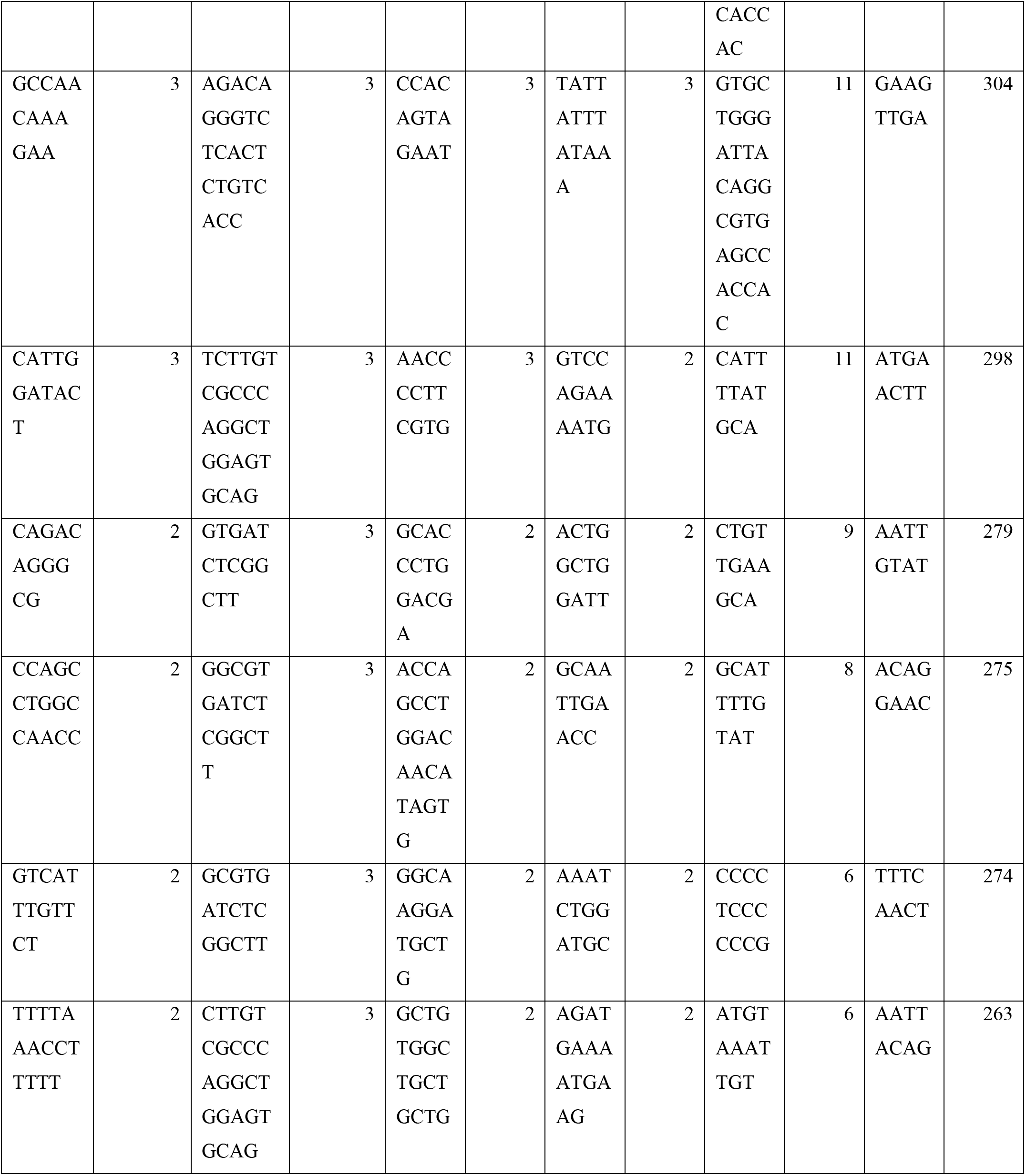
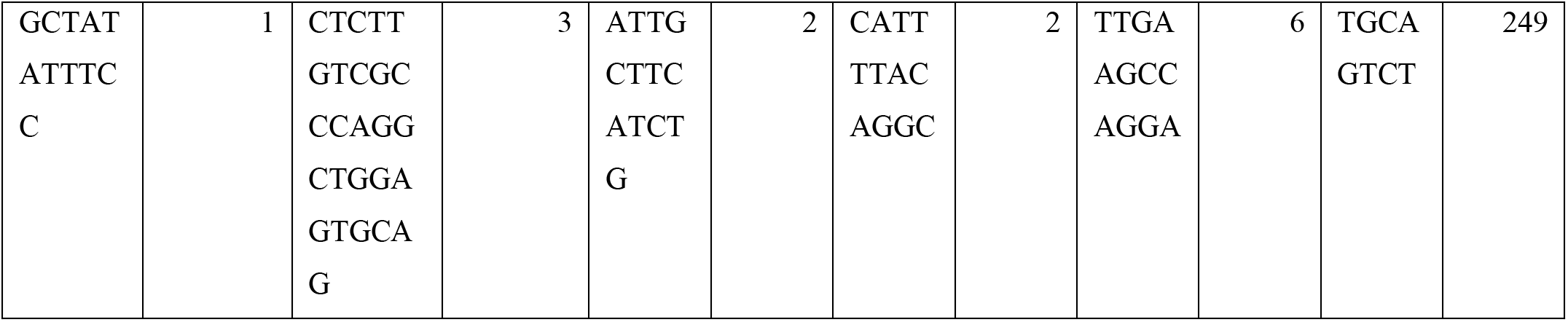
Motifs having the highest hits within each subcellular location

**Table 4.** Motif hits in each subcellular location

|  | # no. of motifs | # no. of hits |
| --- | --- | --- |
| Ribosome | 16 | 32 |
| Cytosol | 32 | 25 |
| ER | 104 | 65 |
| Membrane | 14 | 29 |
| Nucleus | 13 | 95 |
| Exosome | 10 | 1655 |

### Performance of alignment-free methods

#### Performance of CNN based model using one-hot encoded sequence features

A multiscale CNN model was developed using standard python packages – *tensorflow* and *keras*. Training of the model was done using one-hot encoded sequences, each of dimension 2000 × 4.5 fold cross validation was used to evaluate the model without any biasness and preventing overfitting. Due to RAM constraints, we were not able to generate vectors of higher length. The results for the CNN model are attached below-

**Table 5.** Performance of multiscale CNN classifier

| Locations | Sensitivity | Specificity | Accuracy | MCC | F1-score | AUROC |
| --- | --- | --- | --- | --- | --- | --- |
| Ribosome | 0.499 | 0.581 | 0.556 | 0.074 | 0.406 | 0.573 |
| Cytosol | 0.318 | 0.83 | 0.761 | 0.129 | 0.265 | 0.605 |
| ER | 0.247 | 0.837 | 0.77 | 0.072 | 0.198 | 0.583 |
| Membrane | 0.446 | 0.686 | 0.642 | 0.109 | 0.317 | 0.604 |
| Nucleus | 0.804 | 0.325 | 0.656 | 0.14 | 0.764 | 0.596 |
| Exosome | 0.94 | 0.115 | 0.934 | 0.02 | 0.966 | 0.544 |
| **Average** | 0.542 | 0.563 | 0.72 | 0.091 | 0.486 | 0.584 |

#### Performance of machine learning model using composition-based features

Different composition-based features were used to train the ML model and we achieved maximum average AUROC of 0.705 and 0.691 and average MCC of 0.213 and 0.195 respectively in the case of CDK and RDK. Upon combination of these two features, the model performance further improved, giving AUROC in the range of 0.709 and MCC of 0.216.

Training of models were done using standard python packages – Decision Tree, Random Forest Classifier, MLP Classifier, AdaBoost Classifier, Gaussian Naïve Bayes, Quadratic Discriminant Analysis, Gradient Boosting Classifier and Xtreme Gradient Boosting Classifier. Results for all the models are attached in supplementary table. Initially composition-based features were used for training the models and XGBoost classifier had the best performance amongst all the ML models. The metrics for the best performing model is attached below-

**Table 6.** Performance of native XGBoost multilabel classifier

|  | **Sensitivity** | **Specificity** | **Accuracy** | **MCC** | **F1-score** | **AUROC** |
| --- | --- | --- | --- | --- | --- | --- |
| Ribosome | 0.654 | 0.675 | 0.668 | 0.306 | 0.545 | 0.721 |
| Cytosol | 0.648 | 0.649 | 0.649 | 0.208 | 0.334 | 0.703 |
| ER | 0.641 | 0.607 | 0.611 | 0.16 | 0.274 | 0.668 |
| Membrane | 0.67 | 0.669 | 0.669 | 0.27 | 0.431 | 0.732 |
| Nucleus | 0.668 | 0.658 | 0.665 | 0.304 | 0.733 | 0.728 |
| Exosome | 0.546 | 0.731 | 0.548 | 0.048 | 0.706 | 0.706 |
| **Average** | 0.638 | 0.665 | 0.635 | 0.216 | 0.504 | 0.71 |

#### Final model – XGBoost + Motif module

The best performance was obtained on combining the alignment-free XGBoost model with the motif module. Once the prediction is made by the ML model, presence of these motifs was then used to tweak the prediction. If any motif is found in the query sequences, the prediction probability from the ML model is switched to 1. For instance, if a motif unique to ribosome is found within a query sequence, and the prediction probability for that location by the ML model is 0.4, then in the final model it will be assigned a probability of 1. And if no motifs were found for the same query sequence, then the prediction probability assigned by the ML model remains unchanged.

Significant improvement in performance was observed upon implementing this module. The metrics for the ML + Motif module is attached below -

**Table 7.** Performance of XGBoost multilabel classifier combined with the motif module

|  | **Sensitivity** | **Specificity** | **Accuracy** | **MCC** | **F1-score** | **AUROC** |
| --- | --- | --- | --- | --- | --- | --- |
| Ribosome | 0.664 | 0.664 | 0.664 | 0.304 | 0.545 | 0.728 |
| Cytosol | 0.65 | 0.65 | 0.65 | 0.211 | 0.335 | 0.708 |
| ER | 0.657 | 0.656 | 0.656 | 0.205 | 0.304 | 0.727 |
| Membrane | 0.675 | 0.676 | 0.676 | 0.28 | 0.437 | 0.736 |
| Nucleus | 0.671 | 0.671 | 0.671 | 0.319 | 0.738 | 0.736 |
| Exosome | 0.696 | 0.692 | 0.696 | 0.073 | 0.82 | 0.816 |
| **Average** | 0.669 | 0.668 | 0.669 | 0.232 | 0.53 | 0.742 |

#### Comparison of MRSLpred with DM3Loc

Currently, DM3Loc is the only tool that performs multilabel classification out of the box and is developed on the latest version of RNALocate (version 2). We have also used the same non-redundant dataset that is used in DM3Loc. However, the dataset splitting is done differently, maintaining a similar proportion of subcellular locations in each split. Our method manages to achieve a performance similar to DM3Loc. But the major advantage of MRSLpred is that it is computationally very efficient and consumes very less time. Time comparison between both the methods was performed on an iMac (21.5-inch, Late 2015 model, 2.8 GHz Quad-Core Intel Core i5, 8 GB DDR3 RAM and Intel Iris Pro Graphics 6200-536 MB).

**Table 8.** Comparison of MRSLpred with DM3Loc based on time taken

|  | No. of nucleotides | Real time | User time | System time |
| --- | --- | --- | --- | --- |
| DM3Loc | 20 | 0m47.609s | 1m13.874s | 0m2.402s |
|  | 50 | 2m2.065s | 2m48.248s | 0m3.526s |
|  | 100 | 3m28.185s | 5m28.421s | 0m6.338s |
|  | 500 | 13m57.518s | 28m2.371s | 0m42.828s |
|  | No. of nucleotides | Real time | User time | Sys time |
| MRSLpred | 20 | 0m13.659s | 0m3.999s | 0m0.950s |
|  | 50 | 0m22.970s | 0m5.101s | 0m1.020s |
|  | 100 | 0m24.121s | 0m6.965s | 0m0.971s |
|  | 500 | 0m19.649s | 0m20.237s | 0m0.783s |

## Discussion and Conclusion

mRNA localization is a relevant biological process which controls the concentration of mRNA at different location. This, in turn exerts control on the level of protein translation at various locations within the cell [22]. Understanding mRNA subcellular localization will provide a better perspective on how protein synthesis is regulated at the level of mRNA. Spatial localization of mRNA is also gathering huge interest in the field of development biology. Majority of developmental processes rely on cellular polarity generated by mRNA localization to undergo differentiation [23,24]. A lot of research work is focused on understanding these spatial processes which when perturbed can lead to major developmental disorders.

However, in-vitro investigation of mRNA localization is a costly business and labor intensive at the same time. Computational techniques can provide a viable solution to this problem, as they are very fast, inexpensive and reliable. Modern machine learning and deep learning techniques manage to perform well on biological data and are fairly accurate.

A lot of tools have already been developed for mRNA subcellular localization prediction. DM3Loc [1] is a popular tool that deploys multi-head self-attention mechanism with a CNN model. DM3Loc uses a one-hot encoding vector as feature for the CNN model and achieves an AUROC in the range of 0.6981 - 0.7725 (average:0.7415) and a MCC in the range of 0.0736 - 0.3859 (average:0.2696) on their own validation dataset. As of now, only DM3Loc performs multilabel classification whereas all the other tools are multiclass classifiers. Due to its complex architecture, DM3Loc requires very intensive computational power, that includes a dedicated graphics card. Also, for a large number of sequences, one-hot encoding mRNA sequences is a herculean task, as most of the times the RAM crashed due to increasing size of the vector.

MRSLpred achieves AUROC between 0.708 – 0.816 (average:0.742) and MCC between 0.073 – 0.319 (average:0.232) on the validation dataset. MRSLpred uses a much simpler approach combining a XGBoost model with a motif-based module and uses compositional features for the model. Due to this, the method is extremely fast and computationally inexpensive, while achieving comparable performance with the more complicated method - DM3Loc. The time taken by MRSLpred for predicting the location of 500 mRNA sequences is less than 1 minute, whereas for the same dataset DM3Loc takes about 32 minutes.

Sequence-based classification has its own drawbacks, as it tends to lose structural information, which is also known to play an important role in subcellular location. However, in order to use structure-based features for prediction, more computational power may be required. Also, the dataset used in this study had an inherent class imbalance, which would lead to make the model very biased to classes which are represented in larger numbers (Nucleus and Exosome). In the future version of our tool, we expect to obtain a more balanced dataset, where all locations have equal representation.

In this tool, we have managed to develop a multilabel subcellular localization prediction tool, that can accurately identify all the possible subcellular locations that a mRNA could move to. Our final model was based on a XGBoost multilabel classifier that also deploys a motif module to improve our prediction. We have managed to achieve an AUROC 0.708 – 0.816 (average:0.742) and MCC between 0.073 – 0.319 (average:0.232). Due to the simpler architecture of our tool, it is extremely fast and the standalone can be run on very minimal computational power. We believe this will prove to be a useful tool for biologists who work with mRNA localization. MRSLpred is available online at https://webs.iiitd.edu.in/raghava/mrslpred/ and its standalone can be downloaded from https://webs.iiitd.edu.in/raghava/mrslpred/standalone.php. The standalone is also available on GitHub and is accessible at https://github.com/raghavagps/mrslpred.

## Ethics Approval

Not applicable

## Consent to Participate

Not applicable

## Conflict of Publication

Not applicable

## Acknowledgements

Authors are thankful to the Department of Biotechnology (DBT), Department of Science and Technology (DST-INSPIRE) and Council of Scientific & Industrial Research (CSIR) for fellowships and the financial support. Authors are also thankful to Department of Computational Biology, IIITD New Delhi for infrastructure and facilities.

## Authors’ contributions

SC and GPSR collected and processed the datasets. SC and SP implemented the algorithms and developed the prediction models. SC analyzed the results. SC created the back-end of the web server and, SC and NB created the front-end user interface. SC, NB, and GPSR penned the manuscript. GPSR conceived and coordinated the project. All authors have read and approved the final manuscript.

